# Understanding the Neuro-Cognitive Mechanisms of Orthographic Learning in Humans and Baboons: A Comparative Study Using Domain-Specific Mechanistic and Domain-General Connectionist Models

**DOI:** 10.1101/2025.05.16.654419

**Authors:** Janos Pauli, Benjamin Gagl

## Abstract

Script is a key technology for humans, as mastering reading is essential for successful social participation. Hence, understanding the neuro-cognitive mechanisms underpinning the processes of learning to read is highly relevant. Here, we use two orthographic learning datasets from baboons and humans to investigate how they implement visual orthographic representations in a learning task of known and novel letter strings. We use two connectionist models (i.e., CORnet-Z and ResNet-18) and a mechanistic model (i.e., the Speechless Reader model, SLR) to investigate orthographic learning and infer the underlying neuro-cognitive processes. The connectionist models employ neuronally plausible architectures. The SLR versions are transparent neuro-cognitive models of orthographic decision behavior. Central to the SLR implementations are three types of prediction error representations that we use for computational phenotyping (i.e., pixel, letter, and letter-sequence level prediction errors). This approach allows us to infer the underlying representations in orthographic decisions. First, we fit the models and simulate the datasets to compare their performance (i.e., all models see the same sequence of stimuli as humans and baboons). Second, after comparing the model performance, we evaluate how the orthographic decisions have been implemented based on the representations used in the SLR models. We find that the SLR, especially on the trial-wise metrics, outperforms the CNNs in both datasets, with both connectionist models generating behavioral responses without a considerable overlap with individual human or baboon responses. Inspecting the SLR representations, we found that both species implemented the most informative representations that developed from visual to more complex orthographic representations with increased learning. Thus, we show that domain-specific neuro-cognitive mechanistic models are highly valuable in understanding complex behavior and how it is learned across species.

## Introduction

Efficient reading is crucial as low reading skills are associated with low language skills Huettig & Pickering (2019) and a lower socioeconomic status Sirin (2005). The human-developed script representation of meaning and sound is a central part of our lives and a highly relevant stimulus for cognitive science and computer vision (i.e., (Lecun et al., 1998)). Reading research is dominated by two types of models: mechanistic and connectionist models. Mechanistic models provide a transparent, handcrafted implementation of cognitive processes in reading (Coltheart et al., 2001). This line of research was particularly successful in generating effective remediation programs (i.e., phonics; Galuschka et al.,2014) and descriptions of individual differences in reading behavior (i.e., computational phenotypes; Perry et al.,20 In contrast to these domain-specific models, data-driven connectionist models successfully described benchmark effects in reading behavior (Seidenberg & McClelland, 1989). These models implemented connectionist architectures influenced by ideas from the cognitive science of reading, including visual, phonological, and semantic processing. Recent developments have focused on providing deeper insights into the neuro-cognitive processes involved in reading, leveraging connectionist computer vision models based on Convolutional Neural Networks (CNNs). More specifically, recent research has aimed at modeling both behavioral patterns (Hannagan et al., 2021) and the underlying neuro-cognitive mechanisms of reading (Rajalingham et al., 2020). Here, we explore whether the neurocognitive processes of learning how orthographic stimuli are acquired can be captured with a mechanistic approach that models behavior based on neuro-cognitively plausible reading implementations (Gagl et al., 2021; Gagl & Gregorová, 2024). In addition, we compare these models to computer vision models of which variants have been previously used to study learning with orthographic stimuli (i.e., Baboon learning (Hannagan et al., 2014; Linke et al., 2017)).

Central for investigating the learning orthographic stimuli is a baboon learning dataset that describes the behavior of six baboons differentiating between learned and novel letter strings (Grainger et al., 2012). This dataset is interesting as one can compare naive domain-general computer vision models and animals when learning orthographic decisions. Both baboons and computer vision models start the learning process without prior lexical knowledge and script familiarity. Therefore, one can investigate whether the model architecture implements similar processes by comparing their performance. If the model architecture resembles how baboons learn orthographic decisions, one would expect a high alignment between baboon and model behavior. Initial investigations using significantly different architectures (i.e., deep vs. wide) concluded that both models successfully simulated orthographic decision behavior after learning (Hannagan et al., 2014; Linke et al., 2017). Still, both models failed to simulate initial learning as they showed much higher accuracies than the baboons. Note that the task was very easy at the beginning so that baboons could learn to understand the task (i.e., repeated presentation of only a few words). This suggestion was underlined by initial simulations from mechanistic domain-specific models (i.e., variants of the Speechless Reader Model, SLR) showing a much better model fit, especially when emphasizing the initial learning period (Gagl et al., 2021). Thus, at present, three very different model approaches have been used to investigate primate orthographic learning, with only the domain-specific mechanistic model capable of capturing the slow initial learning of baboons. A limitation of all three studies is that each relies on a single model applied to one dataset.

Here, we implement a study that focuses on the development of orthographic decisions as a primitive of reading and visual word recognition. We start by modeling the baboon dataset as a benchmark (Grainger et al., 2012) but extend our investigation to a second dataset based on human orthographic decisions (Eisenhauer et al., 2019). From the modeling standpoint, four approaches are directly compared to each other and against human and baboon behavior, emphasizing the behavior during the initial learning period. We chose two computer vision models using Convolutional Neural Network architectures : (i) CORnet-Z (Kubilius et al., 2018) and (ii) ResNet-18 (He et al., 2016). CORnet-Z is relevant because it is intended to model the hierarchical organization of the ventral visual stream, with four distinct areas corresponding to the retinotopic defined cortical regions in the ventral visual hierarchy: V1, V2, V4, and IT, and likely learns biologically plausible representations (see: (Hannagan et al., 2021; Kubilius et al., 2018)). Thus, the model comprises an architecture capable of simulating orthographic learning behavior. We chose ResNet-18 as it performed well in object recognition tasks (Ibrahim & Shafiq, 2023) and showing as well considerate comparability with human brain activation in the ventral visual stream (Schrimpf et al., 2018).

In contrast to these connectionist models, we included the mechanistic domain-specific Speechless Reader model (Gagl et al., 2024). The inner workings of the model are transparent such that one can derive model-based neuro-cognitive phenotypes for individual participants, allowing one to inspect the individual implementations, i.e., see (Schurr et al., 2024; Schwartenbeck & Friston, 2016). Central to these phenotypes are three types of representations implemented in the model: (i) a visual representation on the pixel level (i.e., described in (Fu & Gagl, 2025; Gagl et al., 2020a)), (ii) an orthographic representation on the letter and (ii) one on the letter-sequence level (both described in Gagl & Gregorová (2024), for similar ideas see Agrawal et al. (2020)). All representations follow the principles of predictive coding (Clark (2013); Price & Devlin (2011); Rao & Ballard (1999)) and implement the efficient representation of sensory information on their respective repvisual word recognition studies (Fu & Gagl, 2025; Gagl et al., resentative levels. Predictive coding is a neuronally plausible theoretical approach. Previous evaluation studies based on neuronal data showed that these representations are implemented during visual word recognition, thus indicating the biological plausibility of these representations on theoretical and empirical grounds (e.g., the findings that prediction errors correlate with EEG-measured brain activation in posterior electrodes from 150-250 ms and EEG-measured brain activation in bilateral occipital cortex and left V4 Gagl et al. (2020a); Fu & Gagl (2025)). Further, the model implements a lexicon that stores all known letter strings, and the output of the model (i.e., learned vs. novel stimuli) is produced by a lexical categorization module (Gagl et al., 2022). The lexical categorization model can simulate the activation pattern of the so-called visual word form area (Cohen et al., 2002; Lerma-Usabiaga et al., 2018; Taylor et al., 2013; Wandell et al., 2012; White et al., 2019) and recently motivated training tasks that result in significantly increased reading skills (Gagl & Gregorová, 2024).

In the present study, testing more realistic architectures of domain-general connectionist models (in contrast to, e.g., LeNet5 in Hannagan et al. (2014)) gives a better chance of modeling initial learning behavior. In addition to domain-general models, we will inspect domain-specific mechanistic models that not only allow us to investigate their performance but also allow the inspection of which type of representation the best fitting models use to solve the task. As metrics, we include standard session-based measures like accuracy on the task and the squared error compared to human/baboon performance. Also, we will rely on more advanced metrics reflecting trial-level differences between models and participants (Geirhos et al., 2020). Thus, we want to use domain-general and domain-specific computational models to describe initial orthographic learning in baboon and human datasets.

## Method Section

### Computational Models

#### Speechless Reader (SLR) Model

The Speechless Reader model (see (Gagl et al., 2024) for all details) is a neurocognitive computational model based on findings from human 2020b, 2022; Gagl & Gregorová, 2024), that can simulate orthographic decisions, which are a primitive of reading. Orthographic decisions are assumed to be solved based only on visual and orthographic letter string information. Therefore, the model involves (i) pixel-level visual and (ii) letter-level orthographic representations (see Gagl et al. (2024) for details). The SLR representations describe sensory information based on efficient neural codes following the principles of predictive coding (Rao & Ballard, 1999; Clark, 2013).

In addition, the SLR includes a memory and a decision module (see Fig. 1). The memory module consists of a list of stored letter strings relevant to the task (i.e., the lexicon that holds all learned items). Before the stimulus presentation, one derives predictions for future events (i.e., the following letter string) on each of the three representation levels based on the learned and stored items in the memory module (see Eq. 1). The predictions implement a mean calculation over the *n* stored items in the lexicon. For the pixel-level representation, the letter strings are transformed into an image, and a pixel-by-pixel mean is calculated over the image (see Fu & Gagl (2025) for more details). The same calculation is implemented on the letter and the letter-sequence level but with a different input matrix. The matrix consists of a slot for each letter on the letter level. For example, assume a lexicon that includes the words *above, about*, and *write*. The probability for the letter *a* at the first position of the word is .66 (*p*(*a*, 1) = (1+1+0)/3 = .66), for *b* at the first position is 0 and *w* at the first position is .33 (*p*(*w*,1) = (0+0+1)/3 = .33). Thus, the letter level prediction error 2 for *above* given this lexicon is 2 (*LPE*)*above* = (1 − .66) + (1 − .66) + (1 − .66) + (1 − .33) + (1 − .66) = 6/3 = 2) and higher at 3 for *write* (*LPE*(*write*) (1 − .33) + (1 −.33) + (1 − .33) + (1 − .33) + (1 − .66) = 9/3 = 3), combining the difference between 1 (i.e., highest error when letter is not expected) minus the prediction (representing how likely the letter is presented at this position) for each letter of the word. For the letter sequence level, the matrix consists of a number of letters minus one (e.g., for House, it would be H, Ho, Hou, and Hous, so that for 5-letter words, the matrix consists of one row with four columns in which the probability of a letter or letter combination is stored). Overall, the predictions integrate the redundant information stored in the lexicon. Thus, the prediction error represents the deviation from the lexicon. This results in representation levels showing the distance from the lexicon, indicating that learned items are near the lexicon and, therefore, have a lower error than the novel items. Note that after the lexicon is used to derive the prediction, it is not further needed to generate the behavioral output.

**Figure 1:**
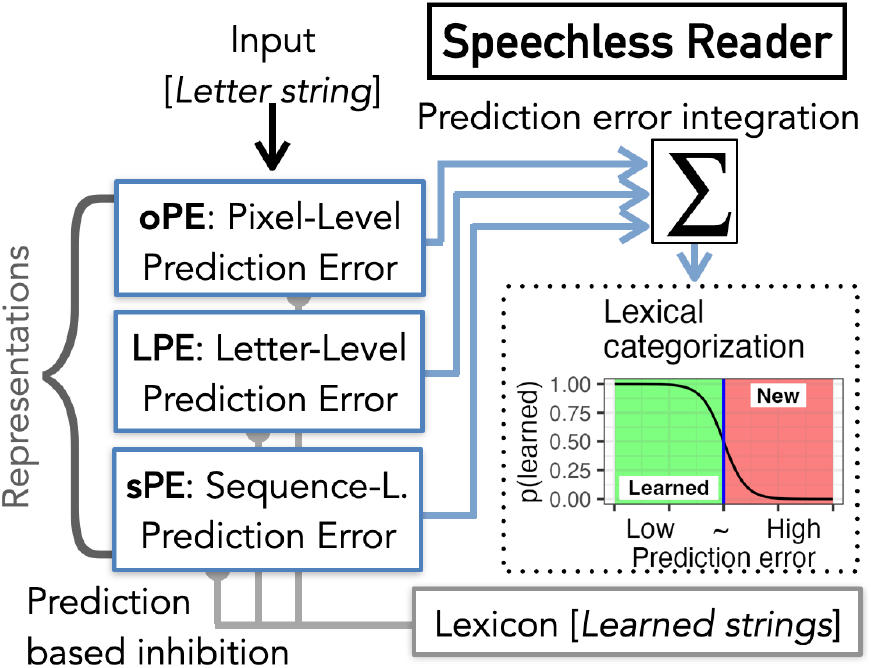
*Speechless Reader Model*, including the lexicon, three types of prediction error representations (i.e., visualorthographic representations, oPE Fu & Gagl (2025); Gagl et al. (2020a); letter-position representations, LPE; lettersequence representations, sPE, Gagl et al. (2024)), and a lexical categorization module that implements the orthographic decision (Gagl et al. (2022)). Prediction errors are calculated based on the learned letter strings stored in the lexicon (i.e., the memory module). The resulting representations are summed into a word-likeness estimate (Gagl et al. (2022)), which is then used as the sole basis for the behavioral output. The output of the categorization process is binary, indicating if the input is considered learned or novel. Note that the model does not implement lexical access as the lexicon only influences the prediction before stimulus presentation.

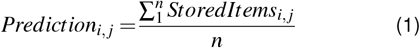

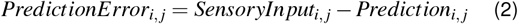

In the next step, the prediction and the sensory input are integrated by subtraction to focus processing on the nonredundant parts of the sensory input. This operation results in a prediction error representation. The SLR first aggregates the prediction error matrices for each representational level (i.e., pixel-, letter- and letter sequence level). These raw prediction error values are then normalized to the same scale to extract a word-likeness measure that consists of one or multiple prediction error representations. Note that the differences in representations combined in the word-likeness measure determine the model variant (i.e., see below). The final step of the model involves a lexical categorization process, which relies solely on the word-likeness estimate. A decision boundary is applied to classify letter strings into two categories: ‘learned’ or ‘novel’. The position of this boundary is the only free parameter adjusted during the model fitting process (i.e., ranges from .05 and ended at .95 in steps of .05). So that the SLR categorizes letter strings with a low word-likeness score (i.e., those that are most distant from the lexicon) as novel items. In contrast, those with a high word-likeness score are classified as learned items.

We adapted the items in the lexicon and the decision boundary for seven model variants that differed in the representations that comprised the word-likeness measure to implement model learning. Letter Strings that were correctly answered more than 80% of the time were added to the lexicon, so the lexicon is likely different for each session and participant. The decision boundary was tested from .05 and ended at .95 in steps of .05 for the word-likeness measures based on the prediction error representations that are normalized between 0-1 (i.e., see schematic description in Fig. 1 lexical categorization, blue line). The boundary allows us to adapt to changing lexicon entries. For example, at the beginning of both tasks, humans and baboons learned only a small set of items. Thus, the prediction error values differ significantly from those observed later in the experiment, where many more items are stored in the lexicon. Therefore, only this one free parameter of the model is needed to account for the very different lexicon setups. Note that we implemented this for all seven model variants. The SLR can be adjusted to determine which representations will be included in the word likeness measure: the model variants. One prediction error alone is sufficient (e.g., a model variant using only the oPE as the word-likeness measure). Still, we implement variants that include all three and all combinations with two prediction errors. Thus resulting in a combined seven model variants determined by all possible combinations of prediction errors (i.e., see Gagl et al. (2024) for details).

We implemented two approaches: One combining the bestfitting and one combining the best-performing models. The best-fitting approach combined the model variants that had a performance most similar to the behavior of the individual participant. The best-performing approach combined the models that had the best performance (i.e., the highest accuracy) at a given data point. The best-fitting approach allows the inspection of the representation likely implemented by the individual reader, and the best-performing approach allows to investigation of the capabilities of the model independent of the performance of the participants.

#### Computer Vision Models and model training

For both models (i.e., CORnet-Z (Kubilius et al., 2018) and ResNet-18 (He et al., 2016)), we set the output layer size to 2, using cross-entropy as our loss function to simulate the binary output in the orthographic decision task (i.e., familiar or novel stimuli). We used the PyTorch (Paszke et al., 2019) library to implement model training. For both datasets, we randomly initialized one network of both computer vision models for each human and each baboon (i.e., six CORnet and ResNet models for baboons and 37 for humans). We set the learning rate to 0.0001 after testing rates from 0.01 to 0.00001 and select the Adam optimizer (Kingma & Ba, 2014). For the model training for the baboon and human data, we transformed all letter strings into grayscale images (width and height of 228 pixels) with a white background and black font. Arial font with a print size of 64 was used for the training set, and for the validation, the same stimuli were presented in Times New Roman, although we only used a validation set for the human data.

To account for previous visual exposure in baboons, we use model instances pre-trained on Imagenet (Deng et al., 2009) and freeze the early layers “conv1”, “bn1” and “layer1” for the ResNet, and the convolutional layers “V1” and “V2” for the CORnet. Human participants were expert readers; thus, we implemented pretrained models that account for the potential transfer effects of literacy on the pseudoword learning task. 37 pre-trained CORnet and Resnet-18 models, one for each human based on a binary classification task (i.e., lexical decision; word or pseudoword) using a dataset with 1,074 5-letter German words and 1,074 5-letter pseudowords. Again, the same early layers are frozen (see above). This way, we match the language input of our pre-trained model and the language of the generated pseudowords from (Eisenhauer et al., 2019)

For the baboon dataset, we trained all models for one epoch, using a pre-trained model instance with froozen early layers. Thus, in each epoch, we trained the model on the same exact stimuli that the baboons saw. This way, we directly mirrored the experimental setup. We calculated the overall accuracy and collected item-level predictions. We did not apply any transformations or augmentations. Here, we do not emphasize the generalization performance of the model since we are solely interested in the learning behavior; see (Hannagan et al., 2014; Linke et al., 2017) for a comparable approach, not implementing a test set.

For the human learning data, we finetuned the pre-trained models on all four epochs of the training dataset. After each we validated the model performance on the respective experimental session using the validation stimuli implementing a different font (i.e., Times New Roman vs. Arial). First, the model is validated on the data from session one after being finetuned on epoch 1. Then, the same is implemented for session two, and so on. Such that we can evaluate the model performance so that it is comparable to the experiment see A1 for training model performance when implementing the same procedure as for baboons). We adopted this validation strategy because the human dataset is relatively small (*N* = 960; *n* = 240 per session), and training on only a subset (e.g., *n* =240) would have limited the reliability of the results.

#### Model evaluation

We modeled performance on the level of session and individual trials for each participant and model. Session-level performance is based on mean accuracy in the task (i.e., is the model correct in the categorization behavior) and the Mean Squared Error (MSE) as compared to the participants (i.e., measures session-wise performance similarity between each computational model and the corresponding subject). The trail-level similarity between the model and participant responses was based on the Error Consistency Metric (see Geirhos et al. (2020) for details). The alignment between two observers, *i, j* (models, participants (baboons or humans), or both), on the same set of stimuli trials *n* can be assessed by measuring the overlap in their response errors (correct or incorrect responses). Since we simulate all our computational models on the identical stimuli set from the behavioral experiments, we can compare item-level responses. For example, we can calculate the observed decision overlap for two different observers, one being a model and the second being a participant. Also, one can compare two participants or two models on this metric. Here, we focus on the participants vs. model overlap and, for comparison, the overlap between participants (within humans and within Baboons).

To properly compare this metric between different participants and models, (Geirhos et al., 2020) suggest plotting it against the expected error overlap. The expected error overlap is based on the accuracy scores of the two observers that are being compared. The metric indicates the probability of making the same decision by chance. The higher the accuracy of the two models, the more likely it is that they made the same decisions by chance. Error Consistency is measured by Cohens Kappa κ. It is based on the observed overlap and the expected overlap by chance. κ > 0 indicates decision consistency above chance, κ = 0 indicates consistency purely by chance, and κ < 0 indicates inverse consistencies, meaning that most likely inverse decision strategies were implemented. If we observe a high consistency in decisions within the two observers, we can assume that they likely share similar internal representations that helped make that decision (McHugh, 2012). Likewise, if we observe consistencies at the chance level, we can assume that the representational systems do not align.

#### Baboon Experiment

For a detailed description of the dataset and task, see Grainger et al. (2012). In short, the dataset consists of item-level accuracy data from over 300,000 orthographic decisions made by six baboons in an orthographic categorization task. The task required determining whether a sequence of letters had been presented multiple times (i.e., a “word”) or not (i.e., a “pseudoword”). The baboons received a food reward for each correct response.

#### Stimuli

The to-be-learned letter strings were randomly selected four-letter words in English, while novel letter strings were four-letter artificially generated pseudowords. All novel items consisted of one vowel and three consonants, with their letter combinations (bigrams) drawn from those found in real English words. Each training session for a baboon consisted of 100 trials. Twenty-five trials included new to-be-learned items, another twenty-five trials randomly presented already-learned items, and the remaining 50 trials consisted of novel pseudoword items. A word item was considered learned when the classification accuracy for it exceeded 80%. The total number of trials varied across baboons (i.e., from 43,000 to 61,100). For each baboon, there was a balance of 50% words and 50% non-words. Since the number of trials varied between baboons, we standardized all baboon datasets by truncating the learning data to the same length (i.e., 43,000).

For model training, we aimed to mirror the experimental setup for each baboon as closely as possible while staying comparable to previous studies that also modeled the learning trajectories in the baboon dataset Hannagan et al. (2014); Linke et al. (2017). Hence, we used the exact same stimuli sequence of the baboon experiments to train the models. We set the batch size to 100, which is one experimental session for a baboon. To make the learning setup comparable with the baboons, we trained the model for one epoch only, as the baboons also did the training just once.

#### Human Experiment

For a detailed description of the dataset, see Eisenhauer et al. (2019). Thirty-seven German-native-speaking human sub-473 jects were familiarized in four sessions with 60 pseudowords (e.g., “Zü nse” or “Posel”), presented twice each session, embedded in 480 unique distractor pseudowords (e.g., “Vedte” or “Lä her”). The participants familiarized themselves with the 60 target pseudowords in each session by reading them aloud once, returning high reading accuracies (a mean error rate of 0.7 % across all sessions). Afterward, the participants did a computerized (old/new) orthographic decision task in which each session contained 240 trials (twice the 60 familiar target pseudowords mixed with 120 novel distractor pseudowords). The participants indicated by button press whether the presented pseudoword was an old, already familiar pseudoword or a novel distractor pseudoword.

#### Stimuli

All non-words were generated with the Wuggy software (Keuleers & Brysbaert (2010). Target and distractor pseudowords shared similar orthographic structures as indexed by the Orthographic Levenshtein Distance 20 (OLD20 Yarkoni et al. (2008); Learned pseudowords, *M* = 1.717; *SD* = 0.026), Novel pseudowords; *M* = 1.743 *SD* = 0.027). The high similarity between the target and distractor pseudowords indicates that the task difficulty on the level of orthography is high. Further, no explicit but potentially implicit semantic information was associated with the pseudowords (e.g., see Gatti et al. (2023)).

## Results

### Baboon Data

Baboon and model accuracy data shows the expected increase in performance with the sessions (see Fig. 2, **A**). Baboons started to increase their performance after about 10 sessions. Both SLR approaches showed better task performance much earlier, showing a performance of about 60% for the best fitting and about 70% correct for the best performing SLR approach. In contrast, both computer vision models showed nearly perfect performance, thus showing a very different model behavior as indicated by the large MSE (see Fig. 2, **B**). All models showed similar accuracies after about 10.000 trials. Yet, overall, the lowest means squared error (see Table 1) and highest trial-level error consistency is found for the best-fit SLR model (see Fig. 2,**C** and Table 2). Notably, ResNet had much lower error consistency with baboon responses than both SLR variants but a still higher kappa when compared to the between-baboon consistency. Overall, the CORnet showed a similar but still a lower kappa as the best-performance SLR (see figure 2, C and table 2).

**Table 1:** Mean Squared Error combined over all sessions.

| Model | MSE |  |
| --- | --- | --- |
|  | Baboon | Human |
| Best performance SLR Model | 0.038 | 0.008 |
| Best fit SLR Model | 0.018 | 0.008 |
| ResNet Simulations | 0.056 | 0.067 |
| CORnet Simulations | 0.019 | 0.116 |

**Table 2:** Prediction Error distribution of the Best-Fit SLR for Humans and Baboons in % and the mean differences between prediction errors for learned and novel letter strings. Larger differences (i.e., lower Δ) indicate that the prediction error representation is more informative for lexical categorization. Also, mean error consistency values for each model and session are presented.

| Source | Session | Frequencies | | | Differences ($\Delta$) | | | Error Consistency | | | |
| --- | --- | --- | --- | --- | --- | --- | --- | --- | --- | --- | --- |
| | | oPE | LPE | sPE | $\Delta$ oPE | $\Delta$ LPE | $\Delta$ sPE | BFS | BPS | RN | CN |
| Humans | Session 1 | 18.9% | 97.3% | 91.9% | -0.701 | -1.105 | -1.019 | 0.432 | 0.176 | 0.142 | 0.068 |
|  | Session 2 | 5.4% | 97.3% | 97.3% | -0.483 | -0.928 | -0.949 | 0.353 | 0.169 | 0.010 | -0.012 |
|  | Session 3 | 0% | 100% | 100% | -0.520 | -0.901 | -1.108 | 0.300 | 0.198 | -0.012 | -0.016 |
|  | Session 4 | 0% | 100% | 100% | -0.578 | -1.000 | -1.072 | 0.234 | 0.161 | 0.031 | -0.009 |
| Baboons | Quarter 1 | 59.4% | 65.8% | 67.3% | -0.430 | -0.750 | -0.599 | 0.349 | 0.115 | 0.033 | 0.096 |
|  | Quarter 2 | 47.8% | 81.2% | 78.9% | -0.193 | -0.497 | -0.311 | 0.305 | 0.069 | 0.044 | 0.084 |
|  | Quarter 3 | 33.3% | 87.6% | 91.6% | -0.040 | -0.231 | -0.141 | 0.319 | 0.123 | 0.042 | 0.103 |
|  | Quarter 4 | 30.5% | 97.0% | 88.2% | -0.059 | -0.286 | -0.120 | 0.347 | 0.141 | 0.051 | 0.135 |

**Note.** oPE = Pixel Prediction Error; LPE = Letter Prediction Error; sPE = Letter Sequence Prediction Error; BFS = Best-Fit SLR; BPS = Best-Performance SLR; RN = ResNet; CN = CORnet.

**Figure 2:**
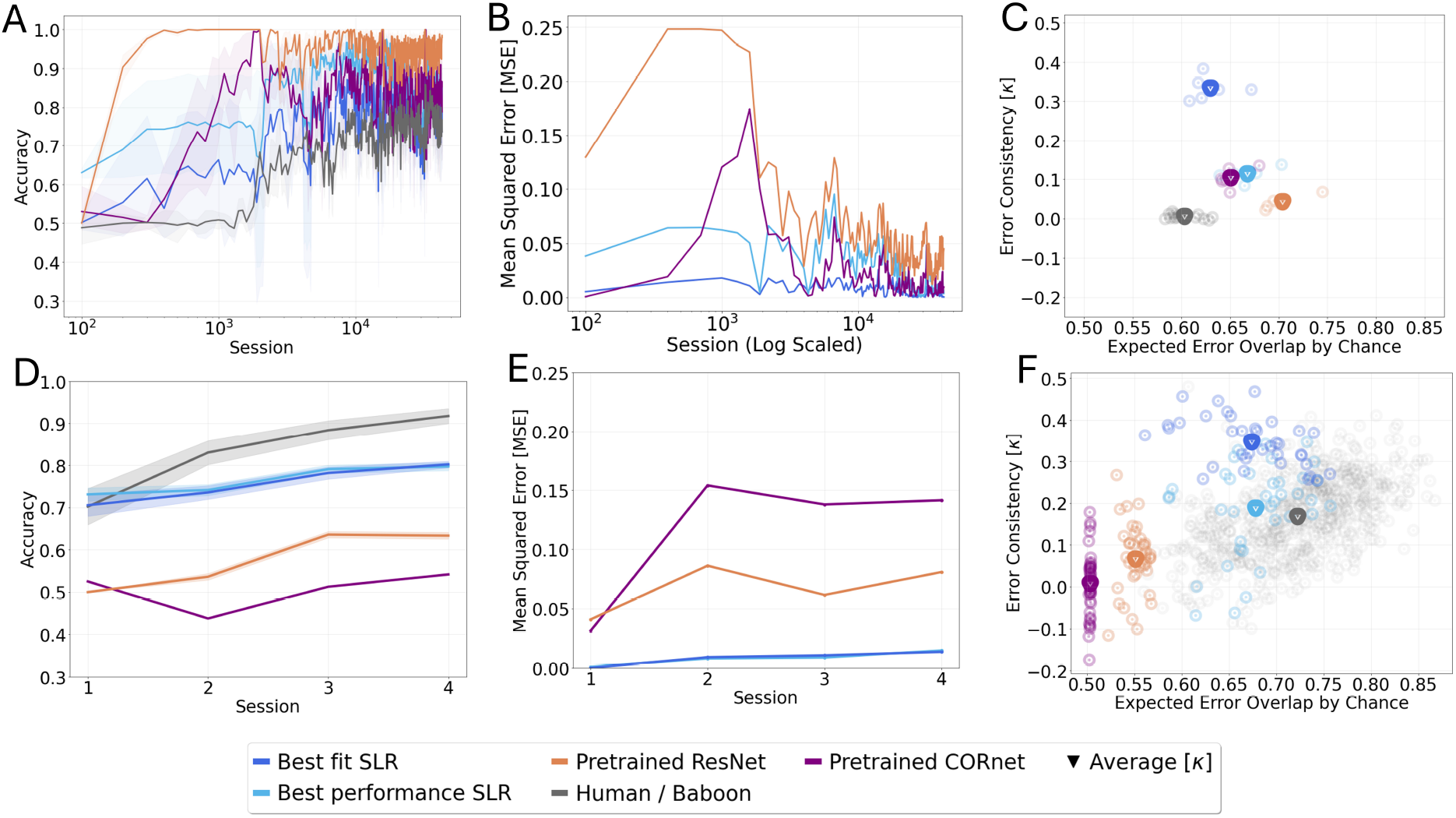
Baboon, human and model performance from the ResNet, CORnet, and two Speechless Reader model implementations (Best-fit and Best-performance variant). (A) Baboon, SLR, and CNN training session-level accuracy, including the 95% confidence intervals. (B) Model fit based on session-level mean squared errors comparing model with baboon performance. (C) Error consistency values (Cohens κ) plotted against the expected error overlap by chance, comparing model and baboon behavior on the level of single trials. Higher κ values indicate stronger item-level behavioral agreement that tends to increase with the expected error overlap (i.e., at high accuracies, the expected overlap is typically higher). The expected error overlap is based on the accuracies of two models (e.g., best-fit SLR and human). The higher the two model accuracies are, the more likely it is that they made the same item-level decisions by chance. In (D), we show session-level accuracy and CNN validation accuracy (E) session-level model fit, and (F) trail-level error consistency for the human dataset.

### Human Data

Mean human and model accuracies and MSE values show the expected increase with learning for all except the COR-net model (see Fig. 2,**D** and Fig. 2,**E**). With incremental learning, the humans increased their performance to above 90% correct, an accuracy unmatched by any model simulation. The CORnet model could not increase the accuracy across sessions, while the ResNet was able to increase the accuracy from 50% to 65%. The best fit and best performing SLR started from about the same level as humans (i.e., at 70% correct), but further learning increase was lower (i.e., at 80% correct after four sessions). Trail-level error consistency overall and separated by session was again highest for the best-fit SLR model simulations and human responses (see Fig. 2,**“F”**) and Table 2). Notably, again, computer vision models had lower error consistency with human responses than both SLR variants. Although ResNet showed a relatively high kappa of 0.142 in the first session, see table 2.

### Neuro-cognitive Phenotypes

Investigating the neuro-cognitive phenotypes provided by the best-fitting SLR model, we find that for both species, all three representations are involved at the beginning of learning (see Table 2). With increasing experience and the involvement of the oPE, visual-pixel-level representation becomes less important for implementing orthographic decisions. Thus, all humans and most baboons rely heavily on the LPE and sPE representations reflecting orthographic processing on the level of letters and letter sequences. This finding is accompanied by increasing differences between learned and novel letter strings in prediction errors, which strongly reduces the oPE representations, ending up with the lowest differences when contrasted to the LPE and sPE representations after learning (see Table 2). Thus, the reduced involvement of the oPE can be explained by a reduction of the differences between learned and novel letter strings, making the representation less informative for the orthographic decision task.

## Discussion

In this study, we offer model-based insights into orthographic learning with a human and baboon dataset. We compare their behavior with model simulations in an orthographic decision task (i.e., discriminating between learned and novel distractor letter strings in a binary decision paradigm). As expected from previous investigations, initial learning periods, especially in the baboon dataset, are highly informative for model comparisons. Additionally, the human dataset, which involved a much more challenging task as stimuli had been more challenging to categorize, allowed good model differentiations since highly similar pseudowords stimuli in the to-be-learned and distractor conditions were presented. Overall, we found that all models could increase the performance, with the exception of the pretrained CORnet in the human dataset. Session-level metrics and trial-level error consistency metrics in both datasets indicated that the mechanistic Speechless Reader (SLR) model approaches described initial learning and overall performance best.

As the precondition of a considerably high model fit was met for the best-fitting SLR model, we could Investigate the neurocognitive phenotypes derived from the model variants using different sets of prediction error representations. The phenotypes indicate a shift from low-level visual pixel-level representations to higher-level orthographic representations with increased learning. Thus, the present study’s focus on learning indicates that orthographic decisions, a primitive of reading, can be accurately described by mechanistic models that also indicate which representations baboons and humans implement to solve the orthographic decision task.

In contrast to previous modeling studies of baboon learning Gagl et al. (2021); Hannagan et al. (2014); Linke et al. (2017), we test more than one model and try to extend the findings to a human pseudoword learning dataset. As expected from previous studies, the critical initial learning data could differentiate between models on the baboon dataset. We found a stark differentiation between the two computer vision models and the mechanistic models, as both the ResNet and CORnet models showed nearly perfect response accuracy after only a few sessions. This performance increase was comparable to previous simulations from computer vision models with vastly different architectures (Hannagan et al. (2014); Linke et al. (2017). This hints that these models are less ideal for investigating such datasets (e.g., see Jacobs & Grainger (1994)).

The finding that the much simpler SLR-based approaches studies are showing that word recognition develops from fragtion of which processes are involved on the individual level models focused on comparisons of data-driven learning archicould simulate both datasets well suggests that we can confidently interpret the representations of the best-fit SLR approach as potentially being implemented by baboons and humans. What we find in both groups, from initial to later learning, is that baboons and humans rely on larger units. At first, pixel-level units are important, but they are reduced in favor of letter and letter-sequence-level representations after initial learning. This finding is in line with more theoretical studies of the inner workings of computer vision models (Agrawal et al. (2020); Hannagan et al. (2021)), as well as neuronal and cognitive theories of efficient reading (e.g., (Coltheart et al., 2001; McCandliss et al., 2003)). Further, a number of developmental mented letter-by-letter-like reading to more efficient reading based on larger multi-letter representations (i.e., see Bijeljac-Babic et al. (2004); Gagl et al. (2015); Huestegge et al. (2009); Schrö ter & Schroeder (2017)) which is conserved to some extent in dyslexic readers (i.e., see (Hawelka et al., 2010; Marinus & de Jong, 2010)) and can reappear after brain injury (e.g., see Gaillard et al. (2006); Pflugshaupt et al. (2009)). Thus, the SLR approach not only provides a model-based transparent way to investigate reading and learning to read (i.e., the best-performing SLR) but also an explicit quantifica(with the best-fit SLR; see also Gagl et al. (2024)).

Using such domain-specific neuro-cognitive mechanistic models can help understand how efficient word recognition is developing, potentially identifying precursors of low reading skills (e.g., see Gagl & Gregorová (2024); Perry et al. (2019)). In addition, each model component can be falsified (e.g., see (Fu & Gagl, 2025)), therefore accelerating the direct progress of how well we understand the neuro-cognitive processes of reading, not only for better identification but potentially also to develop effective training programs (e.g., see Gagl & Gregorová (2024); Galuschka et al. (2014)). Further development and explicit model comparisons could lead to new and vital developments that are not expected from more domain-general tectures (e.g., Gagl et al. (2022); Zorzi et al. (2007)).

Further, using only one free parameter, the SLR approach seems more resilient to data characteristics potentially influencing connectionist models. Inspecting the baboon dataset, we find very different architectures (i.e., LeNET5, Hannagan et al. (2014); one layer reinforcement learning model, Linke et al. (2017); ResNet18 and CORnet, here) showing signs of potential overfitting in initial learning. The baboon task is straightforward at the start, repeatedly showing the same to-be-learned letter strings so the animal can understand the task (i.e., in the first 500 trials, ARI saw only four words with a657 frequency of 7-17 times per session). For connectionist models, the task is inherently implemented in the architecture, as, for example, the input and output modalities are predefined (e.g., the output is defined as binary). Thus, it is unsurprising that the models picked up the to-be-learned words fast as their number is low and the frequency of presentation high. In contrast, in the much smaller human dataset (i.e., in total, 960 stimuli), the task is much more challenging (i.e., to-be-learned and novel pseudowords are highly similar). Hence, the computer vision models have difficulties learning the task. Here, human performance is largest, even unmatched with a pre-trained computer vision model. This finding suggests that humans might use an extended set of multimodal representations, including phonology and semantics (e.g., see Gagl et al. (2024) for a detailed discussion).

What becomes apparent here is that task demands for baboons, humans, and computer vision models are very different, so each group would need specific task implementations. For computer vision and machine learning in general, it is widely known that model training is best with considerable data, testing model performance on left-out test datasets (Brigato & Iocchi, 2021; Huber et al., 2024; Power et al., 2022). Recently, Huber et al. (2024) used a task that allowed the comparison of computer vision and humans on a classic train/test set comparison using pseudo objects with a metalevel structure. Similar situations could also be implemented for pseudoword learning studies. Still, they would create unnatural behavior in this context, as words are letter codes for a specific meaning, not a representation of a meta-level structure. Thus, the advantage of domain-specific models using only a few free parameters is that they are less prone to overfitting and can model datasets with various characteristics within one field.

### Limitations

The current SLR implementation is based on representations accounting only for visual and orthographic aspects of visual word recognition, ignoring phonological and semantic processing influences. Thus, future SLR developments should focus on integrating multimodal representations respecting first phonological processing and potential semantic influences on visual word recognition processes.

We used two versions of the SLR, one implementing a bestfit version. The goal of the best-fit model was to accurately mimic behavior to ensure that the computational phenotypes most reliably represent individual differences. In contrast, the best-performing SLR uses the same lexicon but is not fitted to the individual behavior but uses the model variant with the best performance. Therefore, here, we have no model variant that has an independent lexicon assumption. For component models (i.e., see Fu & Gagl (2025)), we have investigated lexicon structures, showing that the lexicon is likely frequency sorted (i.e., most frequent words first). Here, to achieve lexicon independence, we would need to implement an SLR variant with random sets of to-be-learned letter strings.

## Conclusion

Learning to read is essential for successful social participation. Here, we investigate the initial phase of orthographic learning using domain-general computer vision models (ResNet-18 and CORnet-Z) and domain-specific mechanistic models (Speechless Reader Model variants) based on a baboon and human dataset. We found that these models can all learn visual and orthographic information from letter strings. However, they differ in central aspects, potentially limiting their relevance for the study of orthographic learning (i.e., learning regimes from computer vision models). Central was that the mechanistic models showed respective model fits and provided insights on how humans and primates implement initial learning compared to later skilled processing. Finding that with expertise, we and baboons use larger orthographic units to implement efficient reading, a finding previously shown in studies of children, language learners, and others.

## Appendix

In figure A1 we can see the training performances from COR-net and ResNet, obtained during the fine-tuning on the pseudoword (old/new) task. We fine-tuned each model on the full dataset (*N* = 960). However, to evaluate model performance over time in a way that aligns with the behavioral data, we extracted model outputs session by session. Specifically, we report model performance from epoch 1 on session 1, epoch 2 on session 2, and so forth. This approach allows us to go beyond accuracy metrics and include training data in the calculation of error consistency measures.

Importantly, we could not simply use the entire fine-tuning dataset (i.e., all 960 data points across 4 epochs) for these analyses. This is because the error consistency measures are computed on a session-wise basis and require consistent data structures across sessions—namely, the same number of trials (n = 240) and the same stimulus order as in the human behavioral dataset. These constraints stem directly from the design of the behavioral experiment and ensure that comparisons between model and human data remain valid and interpretable.

**Figure A1:**
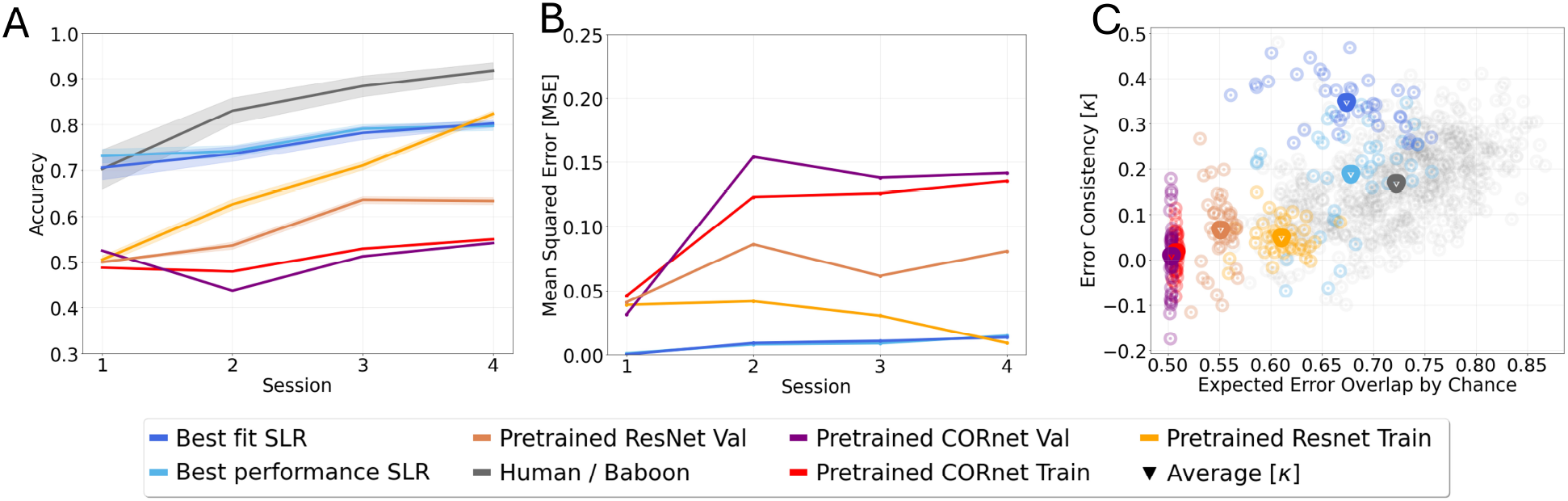
Human and model performance from the ResNet (Train and Validation), CORnet (Train and Validation), and two Speechless Reader model implementations (Best-fit and best-performance variant). In (A), we show session-level accuracy and CNN train and validation accuracy (B) session-level model fit, and (C) trail-level error consistency for the human dataset.

## References

Agrawal, A., Hari, K., & Arun, S. (2020, may). A compositional neural code in high-level visual cortex can explain jumbled word reading. eLife, 9, e54846. Retrieved from https://doi.org/10.7554/eLife.54846 doi: 10.7554/eLife.54846

Bijeljac-Babic, R., Millogo, V., Farioli, F., & Grainger, J. (2004). A developmental investigation of word length effects in reading using a new on-line word identification paradigm. Reading and Writing, 17, 411–431.

Brigato, L., & Iocchi, L. (2021). A close look at deep learning with small data. In 2020 25th international conference on pattern recognition (icpr) (pp. 2490–2497).

Clark, A. (2013). Whatever next? predictive brains, situated agents, and the future of cognitive science. Behavioral and brain sciences, 36(3), 181–204.

Cohen, L., Lehéricy, S., Chochon, F., Lemer, C., Rivaud, S., & Dehaene, S. (2002). Language-specific tuning of visual cortex? functional properties of the visual word form area. Brain, 125(5), 1054–1069.

Coltheart, M., Rastle, K., Perry, C., Langdon, R., & Ziegler, J. (2001). Drc: a dual route cascaded model of visual word recognition and reading aloud. Psychological review, 108(1), 204.

Deng, J., Dong, W., Socher, R., Li, L.-J., Li, K., & Fei-Fei, L. (2009). Imagenet: A large-scale hierarchical image database. In 2009 ieee conference on computer vision and pattern recognition (pp. 248–255).

Eisenhauer, S., Fiebach, C. J., & Gagl, B. (2019). Context-based facilitation in visual word recognition: Evidence for visual and lexical but not pre-lexical contributions. eNeuro, 6(2). Retrieved from https://www.eneuro.org/content/6/2/ENEURO.0321-18.2019 doi: 10.1523/ENEURO.0321-18.2019

Fu, W., & Gagl, B. (2025, Jan). Specifying precision in visual-orthographic prediction error representations for a better understanding of efficient reading. Journal of Cognitive Neuroscience, 1–15. Retrieved from https://doi.org/10.1162/jocna02301 doi: 10.1162/jocna02301

Gagl, B., & Gregorová, K. (2024). Investigating lexical cate-gorization in reading based on joint diagnostic and training approaches for language learners. npj Science of Learning, 9(1), 29.

Gagl, B., Hawelka, S., & Wimmer, H. (2015, apr). On sources of the word length effect in young readers. Scientific Studies of Reading, 19(4), 289–306. Retrieved from https://doi.org/10.1080%2F10888438.2015.1026969 doi: 10.1080/10888438.2015.1026969

Gagl, B., Richlan, F., Ludersdorfer, P., Sassenhagen, J., Eisenhauer, S., Gregorova, K., & Fiebach, C. J. (2022, Jun). The lexical categorization model: A computational model of left ventral occipito-temporal cortex activation in visual word recognition. PLOS Computational Biology, 18(6), e1009995. Retrieved from https://doi.org/10.1371/journal.pcbi.1009995 doi: 10.1371/journal.pcbi.1009995

Gagl, B., Sassenhagen, J., Haan, S., Gregorova, K., Richlan, F., & Fiebach, C. J. (2020a). An orthographic prediction error as the basis for efficient visual word recognition. NeuroImage, 214, 116727.

Gagl, B., Sassenhagen, J., Haan, S., Gregorova, K., Richlan, F., & Fiebach, C. J. (2020b). An orthographic prediction error as the basis for efficient visual word recognition. NeuroImage, 214, 116727. Retrieved from https://www.sciencedirect.com/science/article/pii/S1053811920302147 doi: 10.1016/j.neuroimage.2020.116727

Gagl, B., Weyers, I., Eisenhauer, S., Fiebach, C. J., Colombo, M., Scarf, D., … Mueller, J. L. (2024). Non-human recognition of orthography: How is it implemented and how does it differ from human orthographic processing. bioRxiv. Retrieved from https://www.biorxiv.org/content/early/2024/08/13/2024.06.25.600635 doi: 10.1101/2024.06.25.600635

Gagl, B., Weyers, I., & Mueller, J. L. (2021). Speechless reader model: A neurocognitive model for human reading reveals cognitive underpinnings of baboon lexical decision behavior. In Proceedings of the annual meeting of the cognitive science society (Vol. 43).

Gaillard, R., Naccache, L., Pinel, P., Clémenceau, S., Volle, E., Hasboun, D., … Cohen, L. (2006, Apr). Direct intracranial, fmri, and lesion evidence for the causal role of left inferotemporal cortex in reading. Neuron, 50(2), 191–204. Retrieved from https://doi.org/10.1016/j.neuron.2006.03.031 doi: 10.1016/j.neuron.2006.03.031

Galuschka, K., Ise, E., Krick, K., & Schulte-Körne, G. (2014). Effectiveness of treatment approaches for children and adolescents with reading disabilities: A meta-analysis of randomized controlled trials. PloS one, 9(2), e89900.

Gatti, D., Marelli, M., & Rinaldi, L. (2023). Out-of-vocabulary but not meaningless: Evidence for semantic-priming effects in pseudoword processing. Journal of Experimental Psychology: General, 152(3), 851.

Geirhos, R., Meding, K., & Wichmann, F. A. (2020). Beyond accuracy: quantifying trial-by-trial behaviour of cnns and humans by measuring error consistency. CoRR, abs/2006.16736. Retrieved from https://arxiv.org/abs/2006.16736

Grainger, J., Dufau, S., Montant, M., Ziegler, J. C., & Fagot, J. (2012). Orthographic processing in baboons (papio papio). Science, 336(6078), 245–248. Retrieved from https://www.science.org/doi/abs/10.1126/science.1218152 doi: 10.1126/science.1218152

Hannagan, T., Agrawal, A., Cohen, L., & Dehaene, S. (2021, Nov). Emergence of a compositional neural code for written words: Recycling of a convolutional neural network for reading. Proceedings of the National Academy of Sciences, 118(46). Retrieved from https://doi.org/10.1073/pnas.2104779118 doi: 10.1073/pnas.2104779118

Hannagan, T., Ziegler, J. C., Dufau, S., Fagot, J., & Grainger, J. (2014, Jan). Deep learning of orthographic representations in baboons. PLoS ONE, 9(1), e84843. Retrieved from https://doi.org/10.1371/journal.pone.0084843 doi: 10.1371/journal.pone.0084843

Hawelka, S., Gagl, B., & Wimmer, H. (2010, Jun). A dual-route perspective on eye movements of dyslexic readers. Cognition, 115(3), 367–379. Retrieved from https://doi.org/10.1016/j.cognition.2009.11.004 doi: 10.1016/j.cognition.2009.11.004

He, K., Zhang, X., Ren, S., & Sun, J. (2016, June). Deep residual learning for image recognition. In Proceedings of the ieee conference on computer vision and pattern recognition (cvpr).

Huber, L. S., Mast, F. W., & Wichmann, F. A. (2024). Comparing supervised learning dynamics: Deep neural networks match human data efficiency but show a generalisation lag. Retrieved from https://arxiv.org/abs/2402.09303

Huestegge, L., Radach, R., Corbic, D., & Huestegge, S. M. (2009). Oculomotor and linguistic determinants of reading development: A longitudinal study. Vision research, 49(24), 2948–2959.

Huettig, F., & Pickering, M. J. (2019, Jun). Literacy advantages beyond reading: Prediction of spoken language. Trends in Cognitive Sciences, 23(6), 464–475. Retrieved from https://doi.org/10.1016/j.tics.2019.03.008 doi: 10.1016/j.tics.2019.03.008

Ibrahim, R., & Shafiq, M. O. (2023). Explainable convolutional neural networks: A taxonomy, review, and future directions. ACM Comput. Surv., 55(10), 206. Retrieved from https://doi.org/10.1145/3563691 doi: 10.1145/3563691

Jacobs, A. M., & Grainger, J. (1994). Models of visual word recognition: sampling the state of the art. Journal of Experimental Psychology: Human perception and performance, 20(6), 1311.

Keuleers, E., & Brysbaert, M. (2010). Wuggy: A multilingual pseudoword generator. Behavior research methods, 42, 627–633.

Kingma, D. P., & Ba, J. (2014). Adam: A method for stochastic optimization. arXiv preprint arXiv:1412.6980.

Kubilius, J., Schrimpf, M., Nayebi, A., Bear, D., Yamins, D. L. K., & DiCarlo, J. J. (2018). Cornet: Modeling the neural mechanisms of core object recognition. bioRxiv. Retrieved from https://www.biorxiv.org/content/early/2018/09/04/408385 doi: 10.1101/408385

Lecun, Y., Bottou, L., Bengio, Y., & Haffner, P. (1998). Gradient-based learning applied to document recognition. Proceedings of the IEEE, 86(11), 2278–2324. doi: 10.1109/5.726791

Lerma-Usabiaga, G., Carreiras, M., & Paz-Alonso, P. M. (2018). Converging evidence for functional and structural segregation within the left ventral occipitotemporal cortex in reading. Proceedings of the National Academy of Sciences, 115(42), E9981–E9990.

Linke, M., Brö ker, F., Ramscar, M., & Baayen, H. (2017, Aug). Are baboons learning “orthographic” representations? probably not. PLOS ONE, 12(8), e0183876. Retrieved from https://doi.org/10.1371/journal.pone.0183876 doi: 10.1371/journal.pone.0183876

Marinus, E., & de Jong, P. F. (2010). Variability in the word-reading performance of dyslexic readers: Effects of letter length, phoneme length and digraph presence. Cortex, 46(10), 1259–1271.

McCandliss, B. D., Cohen, L., & Dehaene, S. (2003, Jul). The visual word form area: expertise for reading in the fusiform gyrus. Trends in Cognitive Sciences, 7 (7), 293–299. Retrieved from https://doi.org/10.1016/s1364-6613(03)00134-7 doi: 10.1016/s1364-6613(03)00134-7

McHugh, M. L. (2012). Interrater reliability: the kappa statistic. Biochemia Medica, 276–282. Retrieved from https://doi.org/10.11613/bm.2012.031 doi: 10.11613/bm.2012.031

Paszke, A., Gross, S., Massa, F., Lerer, A., Bradbury, J., Chanan, G., … Chintala, S. (2019). Pytorch: An imperative style, high-performance deep learning library. In H. Wallach, H. Larochelle, A. Beygelzimer, F. d’AlchéBuc, E. Fox, & R. Garnett (Eds.), Advances in neural information processing systems (Vol. 32). Curran Associates, Inc. Retrieved from https://proceedings.neurips.cc/paper files/paper/2019/file/bdbca288fee7f92f2bfa9f7012727740-Paper.pdf

Perry, C., Zorzi, M., & Ziegler, J. C. (2019). Understanding dyslexia through personalized large-scale computational models. Psychological Science, 30(3), 386–395. Retrieved from https://doi.org/10.1177/0956797618823540 (PMID: 30730792) doi: 10.1177/0956797618823540

Pflugshaupt, T., Gutbrod, K., Wurtz, P., von Wartburg, R., Nyffeler, T., de Haan, B., … Mueri, R. M. (2009, Jul). About the role of visual field defects in pure alexia. Brain, 132(7), 1907–1917. Retrieved from https://doi.org/10.1093/brain/awp141 doi: 10.1093/brain/awp141

Power, A., Burda, Y., Edwards, H., Babuschkin, I., & Misra, V. (2022). Grokking: Generalization beyond overfitting on small algorithmic datasets. arXiv preprint arXiv:2201.02177.

Price, C. J., & Devlin, J. T. (2011). The interactive account of ventral occipitotemporal contributions to reading. Trends in cognitive sciences, 15(6), 246–253.

Rajalingham, R., Kar, K., Sanghavi, S., Dehaene, S., & DiCarlo, J. J. (2020). The inferior temporal cortex is a potential cortical precursor of orthographic processing in untrained monkeys. Nature communications, 11(1), 3886.

Rao, R. P. N., & Ballard, D. H. (1999, Jan). Predictive coding in the visual cortex: a functional interpretation of some extra-classical receptive-field effects. Nature Neuroscience, 2(1), 79–87. Retrieved from https://doi.org/10.1038/4580 doi: 10.1038/4580

Schrimpf, M., Kubilius, J., Hong, H., Majaj, N. J., Rajalingham, R., Issa, E. B., … others (2018). Brain-score: Which artificial neural network for object recognition is most brain-like? BioRxiv, 407007.

Schröter, P., & Schroeder, S. (2017, Dec). The developmental lexicon project: A behavioral database to investigate visual word recognition across the lifespan. Behavior Research Methods, 49(6), 2183–2203. Retrieved from https://doi.org/10.3758/s13428-016-0851-9 doi: 10.3758/s13428-016-0851-9

Schurr, R., Reznik, D., Hillman, H., Bhui, R., & Gershman, S. J. (2024). Dynamic computational phenotyping of human cognition. Nature Human Behaviour, 1–15.

Schwartenbeck, P., & Friston, K. (2016). Computational phenotyping in psychiatry: a worked example. eneuro, 3(4).

Seidenberg, M. S., & McClelland, J. L. (1989). A distributed, developmental model of word recognition and naming. Psychological review, 96(4), 523.

Sirin, S. R. (2005, Sep). Socioeconomic status and academic achievement: A meta-analytic review of research. Review of Educational Research, 75(3), 417–453. Retrieved from https://doi.org/10.3102/00346543075003417 doi: 10.3102/00346543075003417

Taylor, J., Rastle, K., & Davis, M. H. (2013). Can cognitive models explain brain activation during word and pseudoword reading? a meta-analysis of 36 neuroimaging studies. Psychological bulletin, 139(4), 766.

Wandell, B. A., Rauschecker, A. M., & Yeatman, J. D. (2012, Jan). Learning to see words. Annual Review of Psychology, 63(1), 31–53. Retrieved from https://doi.org/10.1146/annurev-psych-120710-100434 doi: 10.1146/annurev-psych-120710-100434

White, A. L., Palmer, J., Boynton, G. M., & Yeatman, J. D. (2019). Parallel spatial channels converge at a bottleneck in anterior word-selective cortex. Proceedings of the National Academy of Sciences, 116(20), 10087–10096.

Yarkoni, T., Balota, D., & Yap, M. (2008, Oct). Moving beyond coltheart’s n: A new measure of orthographic similarity. Psychonomic Bulletin amp; Review, 15(5), 971–979. Retrieved from https://doi.org/10.3758/pbr.15.5.971 doi: 10.3758/pbr.15.5.971

Zorzi, M., Perry, C., & Ziegler, J. C. (2007). Nested incremental modeling: The cdp+ model of reading aloud. PsycEXTRA Dataset. Retrieved from https://doi.org/10.1037/e527342012-178 doi: 10.1037/e527342012-178

